# A Cell Painting-Based Tool for the Risk Assessment of Mammary Carcinogens and Endocrine Disruptors

**DOI:** 10.64898/2026.09.04.749408

**Authors:** Rayane Achebouche, Olivier Taboureau

## Abstract

Breast cancer is the most common cancer in women worldwide and chemicals disrupting estrogen or progesterone signaling are recognized as potential risk factors. However, chemicals that alter the mammary gland (MG) development and function remain understudied, highlighting the need for additional research in this area. To address this gap, we investigate the relevance of using high-content imaging assays and more specifically, Cell Painting technology to measure cell morphology perturbation caused by chemical exposure and identify morphological features that could characterize mammary carcinogens (MC) risk factors. Using a dataset of MC and non-mammary carcinogens (Non-MC) with Cell Painting profiles from the JUMP-CP dataset, we retrieved 51 compounds: 28 MC, 23 non-genotoxic Non-MC. We characterized the morphological data by non-linear dimensionality reduction (UMAP) and hierarchical clustering. We, then, developed a Guilt-By-Association (GBA) framework comparing multiple configurations of similarity metrics, risk-score aggregation approaches and features representations. Morphological profiles clustered by mechanism of action rather than carcinogenicity label: genotoxic MC produced strong perturbations in endoplasmic reticulum, mitochondria, nucleus, and RNA compartments, whereas hormonally active compounds were indistinguishable from controls, reflecting the lack of functional steroid hormone receptors in the cell line used. Our best GBA configuration achieved an AUC-ROC of 0.630 and an AUC-PR of 0.696. Applied prospectively to endocrine disruptors chemicals, it prioritized clofentezine, 3-methylpyrazole, resorcinol, 2-tert-butyl-4-methoxyphenol, and thiabendazole as candidates for confirmatory testing. This study provides a transparent, interpretable tool for prioritizing chemicals in mammary carcinogenicity assessment, highlights limitations and clarifies where Cell Painting datasets must be complemented by hormone-sensitive models.

**Plain Language Summary:** Testing chemicals for their potential to cause breast cancer relies on long and costly animal studies. So most chemicals we are exposed to have never been evaluated for such risk. We investigated whether Cell Painting, a microscopy method that captures how chemicals change the cell morphology, could offer a faster, animal-free alternative. Using cell images exposed to chemicals with known mammary carcinogen status, we found that cells respond according to a chemical’s mechanism of action. Chemicals that damage cells directly produced clear and consistent changes. However, chemicals acting through hormones were harder to detect, because the cell type used lack some hormone receptors. By comparing each chemical’s cellular profile to those of known carcinogens, we built a transparent scoring tool that flagged several chemicals for further testing including clofentezine and 3-methylpyrazole. This approach supports the 3Rs by guiding chemicals prioritization before animal testing is undertaken.

## 1. Introduction

Breast cancer is the most common cancer in women worldwide (Bray et al., 2024; American Cancer Society, 2026). Although genetic predisposition factors have been characterized as susceptible to be involved in the disease development, other factors like exposure to environmental chemicals is increasingly recognized as a risk factor (Sun et al., 2017). Epidemiological and experimental evidence implicates a broad range of environmental contaminants such as industrial chemicals, pesticides, plasticisers, or hormonally active substances in mammary carcinogenesis (Rodgers et al., 2018). However, a large proportion of compounds to which human populations are regularly exposed have still not been tested for mammary carcinogenicity. Rodent carcinogenicity bioassays for mammary endpoints are particularly demanding, requiring large numbers of animals over lifetime exposures, which raises both practical and ethical concerns. Developing New Approach Methodologies (NAMs) that exploit existing high-content imaging data therefore offers a concrete opportunity to advance the 3Rs: by prioritizing chemicals *in silico* for targeted confirmatory testing, such approaches can reduce and ultimately help replace routine animal use in mammary carcinogenicity screening. This results in a gap between the breadth of the chemical space and the available toxicological knowledge. That is why it is necessary to use computational and high-throughput predictive approaches to prioritize compounds for further evaluation.

For carcinogenicity prediction, Quantitative structure–activity relationship (QSAR) models are the most established computational framework (Chung et al., 2023; Toma et al., 2020). These machine learning models can learn statistical associations between structural features and biological endpoints by encoding molecular structure as descriptors or binary fingerprints. Applied to mammary carcinogenicity, QSAR approaches benefit from curated chemical lists that distinguish mammary carcinogens (MC) from compounds lacking evidence of mammary tumour induction (Non-MC). The MC class presents a wide range of structural diversity, including polycyclic aromatic hydrocarbons, endocrine disruptors and reactive electrophiles. No single structural representation can account for this diversity of mechanisms, which limits what descriptors alone can resolve (Borrel and Rudel, 2022). Therefore, a complementary source of information reporting on the biological response of cells rather than the chemistry of the molecule is needed.

High-throughput Phenotypic Profiling (HTPP) techniques, such as Cell Painting assays, provide a complementary phenotypic dimension (Bray et al., 2016). These assays capture the overall biological response of cells to a chemical perturbation by quantifying hundreds of morphological features across multiple cellular compartments. They, thus, provide functional information that structural descriptors alone cannot deliver. The JUMP-CP dataset (cpg0016) is one of the largest public resources dedicated to Cell Painting. It provides morphological profiles for thousands of compounds, measured from multiple imaging sources under harmonized experimental conditions (Weisbart et al., 2024). This resource enables morphological analyses and the construction of models based on phenotypic similarity to predict mammary carcinogenicity.

We build on a simple assumption of phenotypic similarity: a compound whose Cell Painting profile closely looks like to a known MC could share its biological effects, regardless of its structural class. This assumption makes the morphological space directly interpretable, since the way compounds group within it should reflect shared mechanisms rather than shared chemistry. It also provides a basis for inference, since we can assess the risk of a compound that induces comparable morphological responses of MCs or Non-MCs based on their similarity. This second, inferential use of similarity is formalized by the “Guilt-By-Association” (GBA) framework, which scores a query compound according to its resemblance to annotated references. requires no classifier training and is inherently interpretable, making it particularly interesting for regulatory screening applications (Lee et al., 2011; Schneidewind et al., 2020). In this study, we assess whether Cell Painting morphology can support mammary carcinogenicity risk assessment through phenotypic similarity. Using the mammary carcinogenicity dataset of Kay and Rudel (2024) mapped on JUMP-CP profiles, we first characterize the structure of the morphological data by non-linear dimensionality reduction (UMAP) and hierarchical clustering, and relate the observed groupings to compound class and mechanism of action. We then develop a GBA framework that systematically compares five similarity metrics, three risk-score aggregation strategies, and three feature representations, evaluated by leave-one-out cross-validation (LOOCV). Finally, we apply the best-performing configuration to endocrine-disrupting compounds and suspected endocrine-disrupting compounds at the EU level, from the EDLists, as a prospective prioritization exercise. We provide a balanced evaluation of the potential and limitations of Cell Painting for mammary carcinogenicity assessment by distinguishing exploratory characterization of the morphological landscape from similarity-based prediction, and by interpreting the results within the experimental context of the JUMP-CP dataset.

## 2. Material and Methods

### 2.1 Chemical Data

In this study we gathered a primary chemical dataset derived from the compendium assembled by Kay and Rudel (Kay et al., 2024), which curates experimental evidence of mammary tumour induction from rodent carcinogenicity studies. After removal of exposures lacking valid InChIKey identifiers, the dataset comprised 1,105 compounds: 276 mammary carcinogens (MC) and 829 putative non-mammary carcinogens (Non-MC). Each entry was annotated with genotoxicity classification and hormonal activity summary.

We used a second dataset for prospective risk assessment. The dataset was made of three endocrine disruptor candidate lists from the EDLists database (The Danish Environmental Protection Agency, 2026). This resource is a centralized repository compiled by participating European national authorities to track substances with potential or identified endocrine-disrupting properties. Substances are organized in different sublists corresponding to different levels of regulatory concern. These sublists are named as List I, List II, and List III (respectively for *Substances identified as endocrine disruptors at EU level*; *Substances under evaluation for endocrine disruption under an EU legislation*; *Substances considered, by the evaluating National Authority, to have endocrine disrupting properties*). We retrieved in tabular format and concatenated into a single dataset the lists I and II, as list III is the least reliable, contains the least data, and consists solely of national assessments. Each compound was assigned a unique list category corresponding to the lowest-numbered list in which it appeared, so that substances present in multiple lists were classified according to the highest level of concern. Structural identifiers (SMILES and InChIKey) were obtained from the PubChem database using the PubChemPy python package (version 1.0.4). We then eliminated duplicates with the same InChIKey in order to obtain a non-redundant set of structurally annotated endocrine disruptors. Thus, we gathered 169 unique compounds, with 99 in List 1 and 70 in List 2.

### 2.2 Phenotypic Data

High Throughput Phenotypic Profiling (HTPP) data were used to analyze and characterize changes in cells following the exposure to the different chemicals we selected in our datasets. Thus, we gathered Cell Painting morphological profiles from the JUMP-CP dataset (cpg0016). This dataset includes over 100,000 compounds and measures compound-induced phenotypic changes in U2OS osteosarcoma cells across multiple imaging sources using the standardised JUMP-CP protocol, in which compounds are applied at a concentration of 10 µM and U2OS cells are exposed to each perturbation for 48 h. The full details of cell-line provenance, authentication, culture conditions, are described in the consortium’s data-generation reference (Chandrasekaran et al., 2023, 2024). Imaging and profile-extraction parameters (microscope, channels, and feature definitions) likewise follow that protocol. All morphological profiles were generated by the JUMP-Cell Painting Consortium and reused here and we did not culture cells or acquire images ourselves. Morphological profiles are extracted from Cell Painting images following the same standardized Cell Painting protocol. These images are composed of five grayscale fluorescence channels, each selectively labeling one or more key cellular compartments or organelles. Plate-level quality control was ensured through negative controls consisting of DMSO mock-treated wells, with 32 replicates per 384-well plate and 128 replicates per 1,536-well plate. Positive controls comprised eight reference compounds, replicated four times on 384-well plates and sixteen times on 1,536-well plates. Several preprocessing protocols were implemented to obtain the most relevant data for our study. We selected the protocol that produces data with the best separation between carcinogenic and non-carcinogenic categories while minimizing batch effects and biases introduced by the diversity of the laboratories conducting the experiments that generated the data. Thus, JUMP-CP morphological raw profiles were processed through the following sequential steps: (i) well-position correction, (ii) cell-count variance normalization, (iii) median absolute deviation, (iv) inverse normal transformation, (v) feature selection by variance thresholding, and (vi) Harmony batch correction to remove inter-source technical variability (Korsunsky et al., 2019). The processed dataset was generated using the Snakemake workflow from the jump-profiling-recipe github repository (Broad Institute, 2025). This dataset comprised 748 morphological features per observation. Per-compound representative profiles were computed as the median across all available sources, yielding a single 748-dimensional vector per compound. Of the 1,105 compounds in the primary dataset, 28 MC and 126 Non-MC compounds (including 23 non-genotoxic) were identified in JUMP-CP, forming the morphological analysis cohort (154 compounds total). In the same way, from the second dataset, we identified in JUMP-CP : 9 compounds out of the 99 that are characterized as List I and 14 compounds out of the 70 that are characterized as List II. These compounds were used as primary risk assessment applications.

### 2.3 Descriptive Analysis of Morphological Profiles

Prior to predictive scoring, morphological profiles were subjected to descriptive analysis. To ensure interpretability and limit confounding effects, we restricted both descriptive analyses to a specific subset of compounds. We excluded Non-MC compounds for which genotoxic activity evidence was reported in a previous study (Kay et al., 2024): of the 126 Non-MC compounds, 103 genotoxic were removed, leaving 23 non-genotoxic Non-MC compounds. This decision is based on two considerations: on the one hand, the Non-MC class contains significantly fewer compounds than the MC class, meaning that keeping Non-MC genotoxic compounds in the analysis could introduce a mechanistic bias unrelated to mammary carcinogenicity; second, genotoxicity constitutes a distinct biological signal that may artificially influence dimensional clustering and separation, independent of mammary carcinogenic potential.

#### 2.3.1 Uniform manifold approximation and projection (UMAP)

Uniform Manifold Approximation and Projection (UMAP) was applied to the combined set of DMSO control, MC and non-genotoxic Non-MC morphological profiles from JUMP-CP. UMAP is a nonlinear dimension reduction technique that aims to capture both the global and local structure of the data. The python package *umap-learn* in version `0.5.9` was used to compute the projection, and visualization in two dimensions was performed using python packages *matplotlib* version `3.10.6` and *seaborn* version `0.13.2` (McInnes et al., 2018).

#### 2.3.2 Hierarchically-Clustered Heatmap (ClusterMap)

We aggregated morphological profiles by compound using the median of replicates, regardless of the source. Hierarchical clustering was performed and visualised as a clustermap using *seaborn* python package (version `0.13.2`) with complete linkage, correlation distance. Row colors indicate compound class (MC vs. non-genotoxic Non-MC), and column colors indicate Cell Painting features family. This approach allowed us to observe and compare phenotypic effects of compounds.

### 2.4 Risk Assessment by similarity : The Guilt-By-Association (GBA) Framework

To estimate the mammary carcinogenicity potential of uncharacterized chemicals, we implemented a Guilt-By-Association (GBA) approach based on morphological profiling (Cell Painting). The core assumption is that chemicals inducing similar phenotypic changes share similar Mechanisms of Action (MoA).

For a given query compound *X*, we evaluated its risk by comparing its morphological profile to reference sets of known MC and Non-MC. To evaluate the similarity between a query morphological profile “*q”* and a reference set of morphological profiles *“B”*, we computed similarity scores using five distinct metrics to identify the metric best suited to this feature space. These approaches can be broadly categorized into correlation-based metrics and transformed spatial distances. Once computed, these pairwise similarities must be aggregated; therefore, we subsequently evaluated multiple strategies to translate these profile-level similarities into a single compound-level risk score. However, because the morphological profile is high-dimensional, it was necessary to reduce noise in order to obtain reliable similarity calculations. Therefore, similarity scores were calculated using and without various dimension-reduction methods in order to retain the most informative representations of the data.

#### 2.4.1 Correlation and Angular Metrics

These metrics assess the alignment or correlation of the feature vectors and naturally produce bounded scores. Thus, we used *Cosine Similarity*, which computes the dot product of *L*_2_ -normalized vectors. It measures the directional alignment between profiles while ignoring their absolute magnitudes; *Pearson Correlation* which is functionally equivalent to cosine similarity computed on mean-centered vectors; *Spearman Rank Correlation* which is a non-parametric metric that applies Pearson correlation to the relative ranks of the features.

#### 2.4.2 Distance-to-Similarity Transformation

Unlike correlation metrics, spatial distances used here yield unbounded values. To standardize these outputs for comparative evaluation, we apply an inverse transformation to map the spatial distance into a bounded similarity score which is described in Eq. 1.

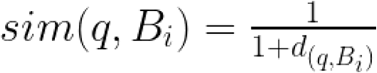

Eq. 1 : **Inverse to Distance Similarity Transformation Formula** This equation shows the conversion of an unbounded spatial distance *d*(*q*, *B_i_*) into a standardized similarity score, where a score of 1 represents identical profiles and values approaching 0 indicate dissimilarity.

We applied this approach to *euclidean distance* to get an *Euclidean Similarity* which remains sensitive to the amplitude of morphological perturbations unlike cosine similarity, and to *mahalanobis distance* to get a *Mahalanobis Similarity* which theoretically accounts for the full inter-feature covariance structure of the reference set. Although we recognize that the low sample-to-feature ratio in our dataset inherently limits the reliability of the empirical covariance estimation we include this metric to investigate whether modeling class-specific biological variance and natural feature correlations provides any exploratory value or detectable signal over unweighted spatial distances.

#### 2.4.3 Risk Score Aggregation Strategies

After having established the mathematical framework for calculating pairwise compound similarities, the next critical step in our GBA pipeline is aggregating these individual profile-to-profile scores into an interpretable risk metric. Because a single query compound is compared against an entire reference matrix of both MC and Non-MC profiles, the method used to pool these similarities significantly impacts the final predictive performance.

We know that carcinogens often operate through highly heterogeneous pathways (e.g., DNA alkylation, oxidative stress, endocrine disruption), leading to distinct morphological signatures. Thus, in order to prevent the mathematical dilution of specific MoA signals, we developed and compared three aggregation strategies, computing a risk score for each. To enable direct comparison of score magnitudes across metrics with differing theoretical ranges, each risk score is divided by the maximum possible value of the corresponding similarity function (1 for Euclidean and Mahalanobis, 2 for correlation-based metrics), yielding a normalized score in (-1, 1) for all strategies.

##### Strategy 1 : Local Neighborhood (Top-*k* Individual Profiles)

This approach assesses the immediate morphological neighborhood of the query compound. To avoid penalizing a query that matches only a rare mechanistic subgroup, we averaged the similarities of only the top-*k* most similar compounds in each class. We set k = 5 as a conservative estimate of the minimum mechanistic subgroup size, consistent with a minimum cluster size of 2 enforced during Dynamic Tree Cut. The individual profile risk score is defined as the difference between the mean similarity of the query to its k nearest MC compounds and the mean similarity to its k nearest Non-MC compounds. The formula is presented in Eq. 2.

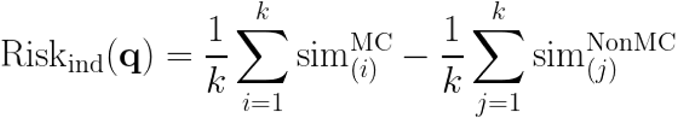

**Eq. 2** : **Top-k Local Strategy Risk Score Formula** With 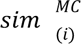 and 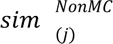 denote the i-th and j-th largest similarity values between the query q and the MC and Non-MC reference profiles, respectively. A positive score indicates a stronger morphological resemblance to MC compounds than to Non-MC compounds.

##### Strategy 2: Mechanistic Subgroups (Cluster Centroids)

MC compounds span multiple mechanisms of action and do not form a single morphological cluster. To explicitly model the MoA subgroups, we performed hierarchical clustering (complete linkage, correlation distance) on both the MC and Non-MC reference sets. We therefore defined each cluster using Dynamic Tree Cut (Langfelder et al., 2007) through *dynamicTreeCut* python package in 0.1.1 version. The parameters used were complete linkage and correlation distance with minClusterSize = 2 and deepSplit = 1. We represented each cluster by its centroid, defined as the arithmetic mean of its member profiles. The risk was then calculated by taking the maximum similarity to any cluster centroid (C). The cluster centroid risk score compares the similarity of the query to its best-matching MC cluster centroid against its best-matching Non-MC cluster centroid as described in the Eq. 3.

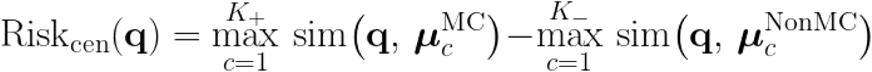

**Eq. 3** : **Cluster Centroid Risk Score Formula** *K*_+_ and *K*_−_ represent the number of MC clusters and Non-MC clusters respectively; µ*_c_* represent the centroid of cluster *c* ;and q represents the query morphological profile. The max operator ensures that a query compound is scored as MC-like if it closely resembles any single MC mechanism of action, irrespective of its dissimilarity to other MC clusters.

##### Strategy 3: Hybrid approach

This hybrid approach combines the top-k and clusters-centroid approaches. This strategy contrasts the maximum MC centroid similarity with the local mean of the k-nearest Non-MC profiles. The resulting risk score which is described in Eq. 4 is designed to maximize sensitivity to specific toxic mechanisms while maintaining a robust, noise-resistant baseline for phenotypic inactivity. This asymmetric formulation encodes the hypothesis that a query compound carries carcinogenicity risk if it looks similar to at least one MC mechanism while its Non-MC likeness is assessed against the full diversity of the Non-MC reference population.

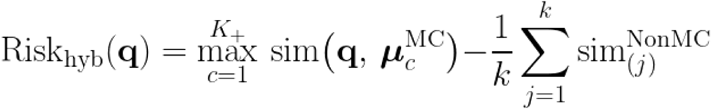

**Eq. 4** : **Hybrid Risk Score Formula** *K*_+_ and *K*_−_ represent the number of MC clusters and Non-MC clusters respectively; µ*_c_* represent the centroid of cluster *c* ; q represents the query morphological profile; and 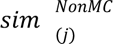 the denotes j-th largest similarity values between the query q and Non-MC reference profiles.

#### 2.4.4 Principal Component Analysis Dimensionality Reduction

To reduce the dimensionality of the data and optimize the performance of our approach, we applied Principal Component Analysis (PCA) (Greenacre et al., 2022) to the combined set of MC and Non-MC morphological profiles. As an unsupervised transformation, PCA does not use the carcinogenicity labels, so the projection basis carries no class information and fitting it on the full reference set therefore does not introduce label leakage into the downstream similarity scoring. Thus, PCA was fitted once on the full combined dataset, and the resulting linear transformation was then applied identically to both classes. We retained the first 7 principal components, which together explain approximately 70% of the total morphological variance. In addition to noise reduction, this transformation provides a geometric advantage specific to distance-based similarity metrics. PCA produces an orthonormal basis ordered by decreasing variance, in which each component encodes the amplitude of morphological perturbation along an independent axis. Euclidean distance computed in this space is consequently closely related to a covariance-normalized metric.

#### 2.4.5 Random Forest Feature Selection

Another approach to reduce dimension was to perform feature selection. To do this, we trained a Random Forest using the following parameters: n_estimators=100, min_samples_split=2 , min_samples_leaf=1, with a train/test ratio of 0.8/0.2. This allowed us to obtain Gini importance scores. We determined the number of the most important features to retain using the elbow method in an automated manner. To do this, we first eliminated the variables with an importance score of exactly zero. Next, we applied to the strictly positive Gini importance scores ranked in decreasing order, the Kneedle algorithm (Satopaa et al., 2011) using *kneed* python package. This method dynamically identifies the point of maximum curvature (the “knee”) on a convex, decreasing curve. Using a standard sensitivity parameter (S=1.0), the algorithm identified 33 most important features to keep when applying this approach on the whole dataset. Because this selection uses the carcinogenicity labels, applying it once to the full dataset and reusing the resulting features across cross-validation folds would allow each held-out compound to influence the features against which it is later scored. To prevent this feature-selection leakage, the Random Forest ranking and Kneedle selection were re-derived independently within every leave-one-out fold, using only the N - 1 reference compounds, so that the held-out compound contributed neither to feature ranking nor to feature selection. We additionally quantified the stability of this selection across folds using the number of selected features and the Jaccard overlap between each fold’s selected set and a reference set derived on the full cohort.

#### 2.4.6 Leave-One-Out Cross-Validation

Having a limited number of annotated compounds, we evaluated the GBA framework using leave-one-out cross-validation (LOOCV). Thus, each compound is used as the query while the remaining N - 1 compounds constitute the reference dataset, yielding 51 independent evaluation folds. LOOCV maximizes the use of available labeled data and is useful for small-sample classification problems (Vabalas et al., 2019). For the full-feature and PCA scenarios, which involve no supervised feature selection, the reference-set construction (clustering and centroid computation) was performed on the N - 1 compounds of each fold. For the Random Forest scenario, feature selection was additionally nested within each fold as described in Section 2.4.5. For PCA and RF scenarios, hierarchical clustering and centroid computation are performed at each fold on the N - 1 reference set of compounds and using the same parameters as described in Section 2.4.3 (complete linkage, correlation distance, minClusterSize = 2, deepSplit = 1). This ensures the query compound does not influence the cluster structure or centroid positions used to compute the risk score keeping the evaluation unbiased and avoid information leakage. Predictions from all 51 folds were pooled, and each performance metric was computed once over this pooled set of held-out predictions (Section 2.4.7).

#### 2.4.7 Performance Metrics and Strategy Selection

Performance of the GBA approach was evaluated using different metrics : AUC-ROC, AUC-PR, MCC, Balanced Accuracy, Sensitivity and Specificity. We prioritize AUC-ROC and AUC-PR as primary metrics for model and strategy comparison, as these threshold-free statistics are insensitive to class prevalence and reflect the ranking quality of the GBA scores. Threshold-dependent metrics (MCC, balanced accuracy, sensitivity, specificity) are reported at threshold = 0 (the natural decision boundary of the signed-difference scoring formula, Section 2.4.3) as secondary, operational descriptors. Thus, for risk assessment applications (Section 3.2.4), ranking-based discrimination (AUC-ROC and AUC-PR) is more relevant than classification threshold dependent metrics. Across all scenarios, the highest values on these metrics were obtained by the cluster-centroid strategy with cosine similarity on the PCA-reduced feature space, which was therefore selected for the prospective scoring of the EDLists compounds.

## 3. Results

### 3.1 Descriptive Analysis

We conducted a two-step descriptive analysis to visualize and understand the structure of the morphological data. The first step allowed observing the data projection in a two-dimensional space, using UMAP, in the aim to identify patterns related to compound class (MC vs. Non-MC) and the data source (imaging center) responsible for generating the profiles. A second step consisted in a hierarchical clustering of the compounds and their morphological features, in the aim to resolve the substructure that a two-dimensional projection cannot display, and to identify which specific features drive the grouping of compounds and how these groupings relate to known mechanisms of action.

#### 3.1.1 Uniform Manifold Approximation and Projection (UMAP)

The UMAP analysis was applied to the combined morphological profiles of MC, Non-MC, and DMSO controls, highlighting compound class and the source from which Cell Painting experiments were conducted.

The projection shows no clear separation between MC and Non-MC profiles in the low-dimensional space. The two classes overlap substantially, suggesting that the morphological signal for mammary carcinogenicity is heterogeneous and not easily separable (Fig. 1).

**Figure 1.**
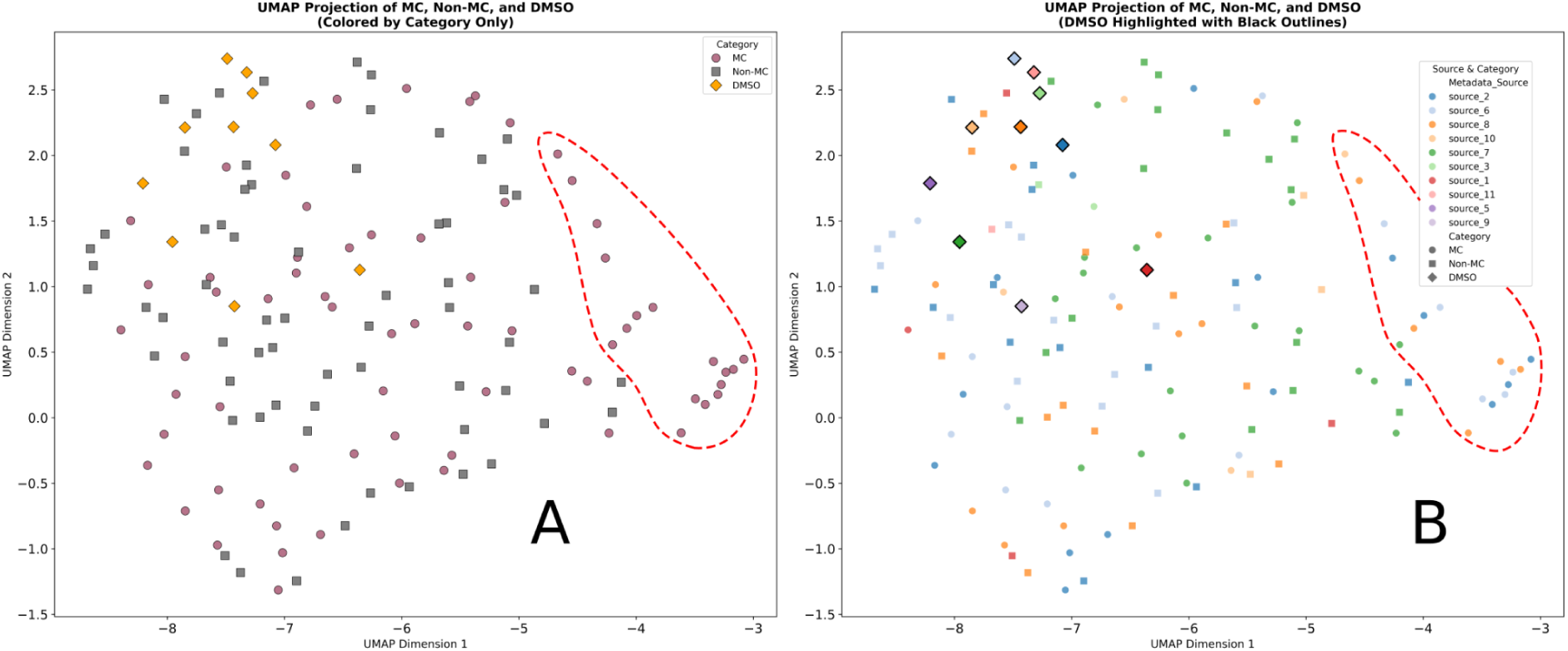
: 2D-UMAP projection of the morphological profiles for cells exposed to MC, Non-MC and DMSO vehicle controls. The dashed red outline marks the compound subset discussed in the text. **(A)** Compounds are colored by category only (MC, circles; Non-MC, squares; DMSO, diamonds). **(B)** Compounds are colored by imaging source (Metadata_Source), with DMSO points additionally outlined in black. Each compound is represented by its median morphological profile aggregated across all available JUMP-CP imaging sources (n biological replicates with n different for each compound). No error bars are shown, as profiles are median-aggregated point estimates.

Although data preprocessing, including normalization and batch correction has been performed using adequate algorithms for Cell Painting data (Korsunsky et al., 2019), a very slight batch effect related to the data source seems to persist and can be observed (Fig. S1). Some sources appear to exhibit profiles that fall within the same region of low-dimensional space, regardless of the nature of the compounds to which the cells are exposed. This source-specific bias could influence the morphological feature values. Furthermore, the median profiles of the DMSO controls for each source were also slightly influenced by this bias; some sources (i.e sources 1; 7; 9) show their DMSO median profile not grouped with other sources. However, we can still observe some trends indicating the presence of different biological signals.

At the bottom-right of the Fig. S1, a cluster of points from carcinogenic compounds originating from different sources converge in the same region. Five molecules are involved: Thiotepa, Folpet, Chlorambucil, Dazomet and Melphalan. They appear to induce a similar phenotypic response, forming a specific cluster of carcinogens.

#### 3.1.2 ClusterMap

Given that UMAP reveals trends showing clusters of morphological profiles exposed to the same types of compounds, it was interesting to see whether a clustering method could validate our first observations. We therefore performed hierarchical clustering combined with a heatmap representation of the median-aggregated morphological profiles of cells exposed to the 51 compounds (28 MCs + 23 non-genotoxic Non-MCs) using complete linkage clustering and correlation distance. This visualization allows us to quickly identify which compounds induce similar morphological profiles and highlight their properties.

##### Dendrogram architecture and global phenotypic heterogeneity

The clustermap reveals two main branches in its compound-dendrogram (Fig. 2). The MC compounds are distributed across both branches, which is consistent with the phenotypic heterogeneity highlighted by the UMAP analysis. Mammary carcinogenicity is not associated with a single, coherent morphological signature in U2OS cells. The structure and organization of the dendrogram branches seem rather to reflect the mechanism of action, and we can observe different mechanistic axes such as direct DNA damage, mitochondrial/oxidative stress, or endocrine disruption.

**Figure 2.**
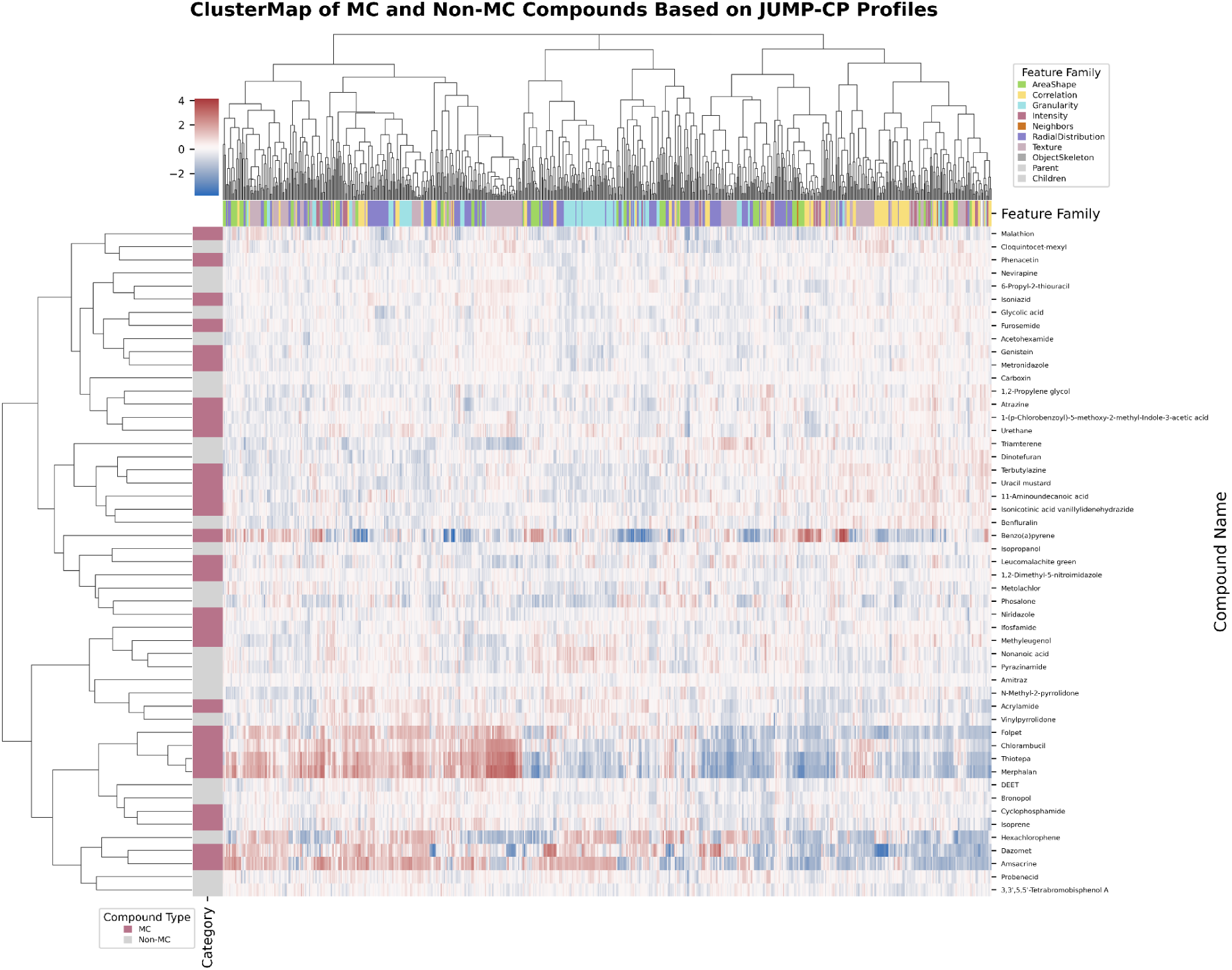
: ClusterMap representation of MC and Non-MC compounds. Compounds are colored according to their Category Type. Features are colored according to the feature family. Each compound is represented by its median morphological profile aggregated across all available JUMP-CP imaging sources (n biological replicates with n different for each compound)). No error bars are shown, as profiles are median-aggregated point estimates.

##### Morphological signatures cluster of Genotoxic, Metabolic, and Cytoskeletal Stress

The main group at the bottom of Figure 2 comprises 13 compounds and contains the highest concentration of MC compounds with strong phenotypic signals. Within the group, we can observe the emergence of subgroups with interesting common attributes (Fig. 2).

The innermost and most compact subcluster is composed of Folpet, Chlorambucil, Thiotepa, and Melphalan which are four MCs with positive genotoxicity. U2OS cells exposed to these compounds exhibit morphological profiles characterized by strong and broad morphological deviations. In particular, significant deviations are observed in Texture Correlation metrics in endoplasmic reticulum (ER), mitochondria (Mito), and RNA channels, with high positive deviations across all four compounds (Cytoplasm_Texture_Correlation_ER_5_01_256 , Cytoplasm_Texture_Correlation_Mito_10_03_256 and Cells_Texture_Correlation_RNA_10_01_256) (Tab. S1). This indicates increased fluorescence homogeneity. One hypothesis is that apoptosis-associated organelle fragmentation, namely endoplasmic reticulum fragmentation (Khaminets et al., 2015) and mitochondrial fission (Chang et al., 2023), breaks normally networked organelles into small, evenly dispersed fragments, smoothing out the local intensity contrasts that the texture feature detects. A similar redistribution of ribosomal RNA could explain the RNA channel signal (Padmanabhan et al., 2012). The Cells_RadialDistribution_RadialCV_mito_tubeness_1 of 16 feature also shows strong positive deviations and indicates disrupted mitochondrial network architecture. This could correspond to mitochondrial fission, which is characteristic of apoptosis induced by DNA damage (Suen et al., 2008).

Despite chemical diversity (Chlorambucil and Melphalan are nitrogen mustards, Thiotepa is an aziridine, and Folpet is a phthalimide fungicide), all four share a genotoxic mechanism. They generate electrophilic species that cross-link or alkylate DNA without requiring metabolic activation (Fu et al., 2012). Folpet reacts with cellular thiols to produce electrophilic species that alkylate DNA and induce strand breaks (Arce et al., 2010). The consistency of this subcluster across the various UMAP imaging sources confirms that the phenotypic signal is robust and does not result from a batch-related artifact.

The second subcluster contains 2 Non-MCs (DEET; Bronopol) and 2 MCs (Cyclophosphamide; Isoprene) with substantially weaker phenotypic effects than the alkylating agent subcluster. Most impacted morphological features involve Actin/Golgi/Plasma membrane (AGP) RadialDistribution (negative deviation of Cytoplasm_RadialDistribution_MeanFrac_AGP_2of4) and DNA-organelle Correlation features (positive deviation of Cells_Correlation_Overlap_AGP_DNA) (Tab. S1). These values indicate a spatial reorganization of the cells. The negative radial distribution shows that actin and Golgi structures have retracted from the wider mid-cytoplasm, while positive overlap scores suggest that either these structures are densely clustered around the nucleus or located directly above it, or that the nucleus is increasing in volume. This signature may reflect a cytoskeletal collapse and cell rounding resulting from acute cellular stress or general cytotoxicity (Desouza et al., 2012).

One Non-MC (Hexachlorophene) and two MC compounds (Dazomet; Amsacrine) compose the third subcluster which is characterized by strong phenotypic effects. Unlike the alkylating agent subcluster, most perturbated morphological features correspond mainly to mitochondrial channels (Nuclei_Intensity_MassDisplacement_Mito, RadialDistribution_FracAtD_Mito_4of4, Cells_Correlation_Overlap_DNA_Mito) as well as Nuclear shape features (Eccentricity features; Zernike moments features) (Tab. S1). These perturbations in U2OS cells reflect a disruption of cellular metabolism and nuclear structural integrity. Mitochondria distribution is being altered, the mitochondrial network is fragmenting and relocalizing which can be a response to metabolic and oxidative stress (Wai and Langer, 2016; Li et al., 2020). At the same time, changes in the nucleus shape indicate a spherization pattern for *Hexachlorophene*, which may also be characteristic of the cell rounding process. Apoptosis induction by cytotoxic compounds has been shown to produce cell rounding concomitant with disappearance of the cytoskeletal network and nuclear fragmentation into rounded nuclear bodies (Jin et al., 2011; Kocgozlu et al., 2010). On the other hand, cells exposed to *Dazomet* (a direct DNA-damaging agent) or *Amsacrine* (a topoisomerase II inhibitor) reveal strong elongation of the nucleus which may be characteristic of genotoxic stress with nuclear membrane destabilization, leading to the leakage of DNA fragments into the cell and often resulting in cell death (Peluso et al., 1998; Boos and Stopper, 2000; Chen et al., 2019; Freyter et al., 2022).

The last subcluster of interest at the base of the 13-compound cluster groups 2 Non-MC : *Probenecid* (an uricosuric) and *3,3’,5,5’-Tetrabromobisphenol A* (a brominated flame retardant). Both produce moderate phenotypic effects without belonging to any of the mechanistically defined groups above. 3,3’,5,5’-Tetrabromobisphenol A is classified as Non-MC in our dataset and as an endocrine disruptor with both E2 and P4 receptor activity. Probenecid is an organic anion transporter (OAT) and pannexin channels inhibitor. Cells exposed to *Probenecid* and *3,3’,5,5’-Tetrabromobisphenol A* show high features deviations for correlation, radial distribution, granularity, and shape features (Cells_Correlation_Manders_ER_DNA, Cells_Correlation_Manders_Mito_DNA, ’Cytoplasm_RadialDistribution_FracAtD_mito_tubeness_2of20, Nuclei_Granularity_11_RNA…) (Tab. S1). This may indicate mitochondrial and endoplasmic reticulum stress. For example, disruptions in radial distribution tubeness metrics reveal that the mitochondrial network is fragmenting, correlation features show the physical dissociation of the ER from the Golgi network and granularity metrics capture the abnormal accumulation of large ER and RNA aggregates around the nucleus (Chandran and Machamer, 2012). This feature profile describes a cell that may shut down its metabolic machinery, suffer nuclear membrane failure, and round up to undergo apoptosis (Balsa et al., 2019).

##### Hormonally active compounds, low phenotypic response and unique profiles

Compounds in the upper branch of the dendrogram (from Malathion to Niridazole) induce slighter phenotypic variations in U2OS cells than the lower branch (Fig. S1). Notably, this upper branch is enriched for compounds known to exhibit estradiol (E2) and/or progesterone (P4) activity, such as Malathion, Atrazine, and Phosalone. The co-clustering of these chemically unrelated compounds in a region of weak morphological deviation is mechanistically coherent. Like other osteosarcoma cell lines, U2OS cells lack functional expression of sex steroid receptors, including estrogen , progesterone, and androgen receptors due to promoter DNA methylation (Dohi et al., 2008; Lillo Osuna et al., 2019). Thus, this steroid receptor deficiency likely explains the lack of phenotypic effects observed for the E2- and P4-active compounds in our dataset.

Like the lower branch, compounds composing the upper branch are very chemically heterogeneous. Some of these compounds show particular phenotypic effects. Indeed, we can observe that *Benzo(a)pyrene* shows strong deviation in morphological profiles compared to other compounds from the upper branch and stands out from the rest. Most perturbed morphological features are associated with DNA channels revealing a very specific phenotypic profile (i.e Nuclei_Granularity_1_DNA, Cytoplasm_RadialDistribution_MeanFrac_DNA_1of4, Cells_Correlation_Overlap_DNA_ER, Cells_Intensity_MassDisplacement_DNA) (Tab. S1). This cell response to the exposition of *Benzo(a)pyrene* may be indicating chromatin fragmentation and formation of micronuclei. This aligns with *Benzo(a)pyrene* mechanism of forming DNA adducts that lead to chromosomal breaks (De Marco Zompit and Stucki, 2021; Liu et al., 2023). The produced morphological profile also reveals that DNA fragments accumulate in the inner perinuclear space and that the nuclear symmetry is preserved unlike what we observed for alkylating/metabolic stress subclusters. This suggests that *Benzo(a)pyrene* induces isolated genotoxic stress and chromosomal fragmentation without triggering the collapse of cytoskeletal and metabolic networks that drive the clustering of lower dendrogram branch subclusters. The rather unique profile of this compound is also reflected in the dendrogram, where it appears relatively isolated compared to the other compounds in the dataset. In the upper branch, two other Non-MC compounds (*Triamterene* and *Carboxin*) are also fairly isolated in the dendrogram, with quite unique signatures. Unlike previous compounds that induce structural failure of U2OS cells, *Triamterene* produces morphological changes characterized by textural disorganization of the ER and mitochondria, loss of cellular polarity, and disrupted organelle spatial organization (Tab. S1). This may reflect a shift into a defense state without leading to fatal outcomes such as nuclear rupture or cell death (Fulda et al., 2010). In the same way, cells exposed to Carboxin produce an even slighter response to chemical stress. Several genotoxic MC compounds also occupy the upper branch: *Phenacetin*, *Isoniazid*, *Furosemide*, *Genistein*, and *Metronidazole*. All are classified as genotoxic in the dataset, yet they produce weak morphological deviations.

Hierarchical clustering confirms and extends the observations from the UMAP projection. The dendrogram groups compounds based on their specific cellular mechanisms of action rather than separating MC from Non-MC compounds. The analysis highlights different mechanistic modes of action. Some genotoxic agents produce strong, coordinated DNA damage response across multiple organelles, some compounds disrupt mitochondrial network through metabolic or oxidative stress, and some compounds with hormonal activity are indistinguishable from negative controls.

This descriptive analysis converges on the fact that mammary carcinogenicity does not correspond to a unique, homogeneous morphological signature in U2OS cells. UMAP Projection and Clustermap analyses show that mechanism of action is the main principle that determines the profile distribution in the morphological space. Genotoxic compounds consistently separate from the rest of the dataset in both visualizations and their reproducibility across sources in the UMAP confirms that this phenotypic signal is biologically meaningful and technically reliable. Hormonally active compounds including MC produce profiles with muted response which is a consequence of steroid receptor deficiency in the osteosarcoma cell lines such as U2OS. This renders the mechanism of endocrine disrupting driven mammary carcinogenesis phenotypically silent here. The dispersion of MC compounds in UMAP space and in dendrogram space is therefore not a detection issue, but rather a direct morphological reflection of the mechanistic heterogeneity of the MC class. Genotoxic initiators, mitochondrial disruptors, and endocrine disruptors exist in distinct, non-overlapping regions of the morphological space. This structure has a direct impact on the predictive modeling potential. Any classifier trained on the entire MC class is confronted to a configuration that fundamentally limits the performance of global binary approaches regardless of feature selection strategy, model choice or even sample size.

### 3.2 Assessment by Similarity

In the aim to propose a predictive model able to detect possible MC compounds, we evaluated 15 strategy-metric combinations in multiple scenarios. First we applied our GBA framework on the full feature space (748 features, 51 compounds). Then, in a second scenario, we reduced the feature space to its first 7 components through PCA. Finally, we reduced the feature space but this time by selecting the most important features through Random Forest modeling.

#### 3.2.1 Full Morphological Features Space

In this scenario, every strategy-metric configuration produced poor performance with a 0.415 highest AUC-ROC value (Tab. S2). This suggests that for the vast majority of query compounds, Non-MC similarities are greater than MC similarities, introducing a bias that shifts all signed difference risk scores toward negative values. Two closely related factors could explain this consistent underperformance. First, the data suffer from the problem of high dimensionality. With more than 700 morphological features for only 51 compounds, the feature space is vastly disproportionate to the sample size. In this high-dimensional context, the mathematical distances between any two profiles blur and converge, thereby depriving distance-based measures of their ability to reliably classify similarities among compounds. Secondly, the morphological features do not separate distinctly the MC and Non-MC classes. Indeed both groups broadly overlap for most of the compounds as shown in the UMAP projection (Section 3.1.1). The discriminative signal may be deteriorated by the accumulation of noise across the 748 features, the two classes are mixed within the same morphological space, and standard similarity measures cannot effectively distinguish them in this context. It is necessary to consider transforming the data in a way that creates a context conducive to evaluating compounds based on their similarities. We, therefore, tested whether reducing the dimensionality of the space could expose a usable signal.

#### 3.2.2 Principal Components Reduced Feature Space

For this scenario we reduced the feature space to its first 7 principal components (PCA), retaining approximately 70% of total morphological variance (Section 2.2). Because PCA is unsupervised, the projection does not use carcinogenicity labels. The first advantage of this method is the noise reduction, but it also gives a geometric advantage by encoding the magnitude of the morphological perturbation along each variance axis which are by definition orthogonal. The euclidean distance in this PCA-reduced space is therefore very similar to a covariance-normalized metric without requiring an explicit estimate of the covariance which is essential with our sample size (N = 51). Performance following this PCA dimension reduction improved markedly relative to the full space, with 6 of the 15 strategy-metric configurations achieving AUC-ROC above 0.5 (Tab. 1). Cluster centroid strategies combined with cosine similarity (AUC-ROC = 0.630, AUC-PR = 0.696) and Pearson similarity (AUC-ROC = 0.607, AUC-PR = 0.676) achieved the highest discrimination. The latter yielding the most favorable threshold-dependent operating point (MCC = 0.236, balanced accuracy = 0.610). Hybrid strategies with Pearson and cosine similarity produced moderate above random results (AUC-ROC = 0.585 and 0.545). All individual profile strategies remained below chance (AUC-ROC = 0.200 - 0.455) and showed that assuming that carcinogenicity is encoded by proximity to any carcinogen is not a good way to tackle the problem. Indeed, the descriptive analysis (Section 3.1) showed that this assumption does not hold, as the MC class is mechanistically heterogeneous and its members are dispersed across several distinct regions of the morphological space rather than forming a single neighborhood. Mahalanobis-based configurations performed poorly across all strategies (AUC-ROC = 0.200 - 0.408), consistent with the unreliable covariance estimation expected at this sample-to-feature ratio. Overall, aggregating similarities at the level of mechanistic subgroups (cluster centroids) rather than individual profiles was necessary to recover a usable signal, in line with the mechanistic heterogeneity of the MC class established by the descriptive analysis.

**Table 1.** : Performance of GBA framework on PCA selected features space for MC and Non-MC non genotoxic compounds set. Performance metrics were computed once over the pooled predictions from all 51 leave-one-out cross-validation folds (one held-out compound per fold). Because the reference set and configuration are fixed, the procedure is deterministic and yields a single value per metric.

| Strategy | Metric | AUC-ROC | AUC-PR | MCC | Bal.Acc | Sensitivity | Specificity |
| --- | --- | --- | --- | --- | --- | --- | --- |
| cluster_centroids | cosine | 0.630 | 0.696 | 0.155 | 0.575 | 0.714 | 0.435 |
| cluster_centroids | pearson | 0.607 | 0.676 | 0.236 | 0.610 | 0.786 | 0.435 |
| cluster_centroids | euclidean | 0.554 | 0.655 | 0.117 | 0.557 | 0.679 | 0.435 |
| cluster_centroids | spearman | 0.443 | 0.561 | -0.128 | 0.438 | 0.571 | 0.304 |
| cluster_centroids | mahalanobis | 0.408 | 0.595 | 0.000 | 0.500 | 1.000 | 0.000 |
| hybrid | pearson | 0.585 | 0.639 | 0.097 | 0.533 | 0.893 | 0.174 |
| hybrid | cosine | 0.545 | 0.629 | 0.148 | 0.559 | 0.857 | 0.261 |
| hybrid | euclidean | 0.528 | 0.665 | -0.116 | 0.448 | 0.679 | 0.217 |
| hybrid | spearman | 0.439 | 0.587 | -0.204 | 0.408 | 0.643 | 0.174 |
| hybrid | mahalanobis | 0.331 | 0.499 | -0.294 | 0.352 | 0.357 | 0.348 |
| individual | euclidean | 0.455 | 0.566 | -0.082 | 0.460 | 0.571 | 0.348 |
| individual | cosine | 0.390 | 0.532 | -0.139 | 0.434 | 0.607 | 0.261 |
| individual | pearson | 0.379 | 0.528 | -0.092 | 0.456 | 0.607 | 0.304 |
| individual | spearman | 0.287 | 0.452 | -0.454 | 0.280 | 0.429 | 0.130 |
| individual | mahalanobis | 0.200 | 0.400 | -0.299 | 0.411 | 0.821 | 0.000 |

Having established that an unsupervised projection partially recovers the biological signal, we next asked whether a supervised reduction, selecting features explicitly for their discriminative importance, could improve on it.

#### 3.2.3 Random Forest Selected Features Space

In this last scenario we assessed a supervised dimensionality reduction based on Random Forest feature selection. Because this selection uses the carcinogenicity labels, it must be performed within cross-validation to avoid information leakage. We therefore embedded the selection in a nested leave-one-out scheme, re-deriving the important features on the N-1 reference compounds at every fold, so that the held-out compound never contributes to choosing the features against which it is subsequently scored. Random Forest selection did not yield a usable signal: performance was at or below chance for nearly all configurations (AUC-ROC = 0.337–0.646; Tab. S3). The single value above 0.5, obtained by the hybrid strategy with Mahalanobis similarity (AUC-ROC = 0.646), is not usable, as its threshold-based operating point is degenerate (sensitivity = 0.000, specificity = 1.000, MCC = 0.000), meaning every compound is assigned to the Non-MC class. All correlation-based configurations, which performed best in the PCA space, remained close to random here (AUC-ROC = 0.441–0.511). To understand this collapse, we examined the stability of the Random Forest selection across the 51 folds. The selection proved highly unstable: the number of features retained by the Kneedle elbow varied substantially from fold to fold (median 24, range 12 - 40), rather than converging on a fixed set (Tab. S4). More importantly, each fold’s selected set shared, on average, only 3.8 features with the 33-feature set obtained from the RF feature selection on the full dataset. This instability indicates that, at this sample-to-feature ratio, the Random Forest importance ranking does not identify a consistent, generalizable subset of discriminative features.

Consequently, the unsupervised PCA projection, which does not depend on labels and is stable by construction, provides the most reliable configuration for the subsequent risk-assessment application.

#### 3.2.4 Assessment of risk compounds based on similarity

To test our risk assessment method on compounds that may have an impact on breast cancer, we applied our method to lists of compounds suspected of causing mammary carcinogenicity. Thus, the optimal GBA configuration identified in Section 3.2.3 was selected and used to screen compounds from the List I and List II of the EDLists. This configuration combines the cluster-centroid strategy with cosine similarity on the PCA-reduced feature space (7 components) and yielded an AUC-ROC of 0.630, an AUC-PR of 0.696, a sensitivity of 0.714, and a specificity of 0.435. This operating point is more balanced than a purely sensitivity-oriented one, while remaining appropriate for a prioritization framework in which compounds flagged as MC-like are directed toward confirmatory experimental testing rather than treated as definitive classifications. Before scoring the compounds at risk, we excluded one compound from List II because it was already present in the previously established MC/Non-MC reference set. Thus, the final evaluation sets comprised 9 and 13 unique compounds for List I and List II, respectively. Each EDList compound-induced morphological profile was scored as a query against the reference set of 51 annotated compounds (28 MC and 23 non-genotoxic Non-MC) using the selected configuration (cluster-centroid strategy, cosine similarity, PCA-reduced space).

In List I, four compounds were classified as MC-like: Clofentezine (0.313), 4-MBC (0.069), Medetomidine (0.032), and 3-BC (0.031). Clofentezine received by far the strongest MC-like score of the entire screen, more than twice that of any other compound. The remaining five compounds were classified as Non-MC-like, with 4-tert-octylphenol (-0.186) and benzyl butyl phthalate (BBP; -0.094) at the lower extreme; bisphenol A (-0.043), dibutyl phthalate (-0.039), and butylparaben (-0.008) completed this group (Tab. S5).

In List II, six compounds were classified as MC-like, led by 3-methylpyrazole (0.127), resorcinol (0.095), and 2-tert-butyl-4-methoxyphenol (BHA; 0.080), followed by thiabendazole (0.049), butylated hydroxytoluene (BHT; 0.041), and propylparaben (0.009). The remaining seven compounds were classified as Non-MC-like, with triclosan (-0.206) and TDCP (-0.144) at the lower extreme (Tab. S6).

Two aspects of these results deserve to be highlighted. First, the three parabens present across the lists all scored close to the decision boundary (propylparaben +0.009, methylparaben -0.002, butylparaben -0.008), rather than receiving divergent scores. This clustering near zero is consistent with a limited morphological response to compounds whose principal activity is receptor-mediated, in a cell line that does not express functional steroid receptors, and indicates low confidence in their classification either way. Secondly, several UV filters and preservatives with primarily hormonal mechanisms (oxybenzone-0.008, avobenzone -0.011, climbazole -0.041) also yielded scores close to or slightly below zero, forming a low-confidence cluster around the threshold.

These scores require careful interpretation given the cellular context in which the morphological profiles were generated. Indeed, we know that the GBA framework was developed and applied using profiles derived from the U2OS cell line (which does not exhibit significant endogenous expression of estrogen, progesterone or androgen receptors), and that the compounds in List I and List II were specifically selected for their confirmed or suspected endocrine-disrupting activity. Consequently, a Non-MC-like or near-zero score should not be interpreted as evidence of low carcinogenic potential, especially for compounds acting primarily through receptor-mediated hormonal disruption. Rather, it may reflect the limited sensitivity of U2OS cell morphology to receptor-mediated endocrine signaling, regardless of the actual biological activity. This is illustrated by oxybenzone, whose near-zero, marginally negative score cannot be reconciled with its reported estrogen-receptor-dependent genotoxicity in breast epithelial cells (Majhi et al., 2020), and most likely reflects the inability of U2OS to register that receptor-mediated mechanism. Conversely, an MC-like score remains informative regardless of the mechanism, since it indicates that the compound induces morphological changes resembling those of known mammary carcinogens via pathways detectable in this system. This may include genotoxic, cytotoxic, or oxidative stress-related mechanisms, or simply side effects of receptor activation via cross-talk signaling.

On this basis, and prioritizing compounds that combine a strong MC-like score with a mechanism plausibly detectable in a receptor-deficient line, Clofentezine, 3-methylpyrazole, resorcinol, BHA, and thiabendazole emerge as the primary candidates for confirmatory testing. The high score of Clofentezine, a known rodent hepatocarcinogen, is notable and makes it the strongest candidate of the screen. 4-MBC also received a positive score, despite being known for its essentially estrogenic mechanism. Since hormonal receptors-mediated effects are not expected to be detectable in U2OS cells, this positive signal could indicate a secondary independent activity, warranting careful attention. Conversely, compounds classified as Non-MC-like whose principal mechanism is receptor-mediated (e.g., the parabens and oxybenzone) cannot be cleared on that basis and must be validated in a hormone-sensitive cell line (e.g., MCF-7 or T47D) before their GBA classification can be considered informative (Nowak et al., 2018). A negative score in U2OS does not currently distinguish a true absence of carcinogenic potential from a failure to detect it.

## 4. Discussion

In this study, we investigated whether the phenotypic response captured by Cell Painting could be used as an indicator of mammary carcinogenicity. Our analyses led us to a key observation showing that morphological profiles are organized according to mechanism of action rather than their carcinogenicity label directly. Based on this observation, we were then able to propose a predictive score to assess the risk of molecules based on the similarity of their morphological profiles to the reference MC and non-MC phenotypic signatures.

In both the UMAP projection and the hierarchical clustering, MC compounds do not form a coherent group. In fact, they are distributed across the genotoxic, mitochondrial/oxidative stress, and hormonally active axes, which also contain non-MC compounds. This observation frames all the results that follow and explains why the overall separation of classes is weak, while local mechanism-aware similarity is informative.

The Cell Painting technique reveals biologically significant and reproducible signatures for compounds that act through direct genotoxic or cytotoxic mechanisms. The subgroup of alkylating agents (Folpet, chlorambucil, thiotepa, melphalan) is the most obvious example: these signatures are consistent across different imaging sources, indicating that the phenotypic signal is robust and not the result of a batch-related artifact. This justifies the use of the Cell Painting technique as a preliminary filter for chemicals whose mode of action leads to obvious morphological changes. This initial morphological assessment is, however, limited by the cellular context in which the JUMP-CP profiles were generated. U2OS osteosarcoma cells do not express functional estrogen, progesterone, or androgen receptors (Lillo Osuna et al., 2019). Thus, compounds that act primarily through receptor-mediated endocrine signaling are phenotypically silent in this system. Hormonally active compounds, including several MCs, are therefore grouped with the inactive controls. As a result, in an assessment based on JUMP-CP profiles, an MC-like score is informative regardless of the mechanism, whereas a Non-MC-like score or one close to zero does not allow one to distinguish between a true absence of carcinogenic potential and an inability to detect a receptor-mediated effect in a receptor-deficient cell line.

Descriptive analysis directly reveals the limitations of a global supervised model. Since the MC class is mechanistically heterogeneous and its members occupy distinct, non-overlapping regions of morphological space, any classifier trained to separate the entire MC class from the entire Non-MC class encounters a bottleneck that limits achievable performance, regardless of feature selection, model choice, or sample size. The GBA framework offers an interesting alternative because it does not attempt a global separation. Indeed, by evaluating a query relative to its closest MC neighbors or to the centroid of the MC cluster that most closely fits it, it exploits the local mechanistic structure. This is consistent with the failure of every strategy-metric configuration in the full 748-feature space (best AUC-ROC 0.415). Reducing the dimensionality with an unsupervised PCA projection recovered a biological signal (best AUC-ROC 0.630, AUC-PR 0.696, for the cluster-centroid strategy with cosine similarity), indicating that the failure in the full space reflected noise accumulation across many correlated features rather than a complete absence of discriminative structure. Supervised Random Forest feature selection, by contrast, showed poor performance, and a stability analysis revealed that the selected feature sets varied substantially from fold to fold and shared only a handful of features with any globally selected set. At this sample-to-feature ratio, no stable, generalizable discriminative subset could be identified, and the unsupervised projection, being label-independent and stable by construction, provided the only reliable configuration. The fact that the signal emerges only when similarities are aggregated over mechanistic subgroups (cluster centroids) rather than individual profiles further reflects the heterogeneity of the MC class. Prediction succeeds better by exploiting local mechanistic structure rather than attempting global class separation.

In the context of risk assessment or regulatory screening, the properties of the GBA approach are just as important as its raw performance. Indeed, this approach requires no trained model, its score corresponds to the signed difference between the similarity to known carcinogens and that to non-carcinogens, and each prediction can be linked to the specific reference compounds and the mechanistic group that determined it. This interpretability is difficult to achieve with “black-box” classifiers and is essential for identifying the relevant molecular characteristics, elucidating the underlying biological mechanisms, and justifying regulatory decisions with full transparency. For a prioritization use case, the quality of the ranking matters more than any single decision threshold, which is why we selected the configuration on threshold-independent metrics (AUC-ROC and AUC-PR). The selected configuration achieves moderate, above-chance discrimination (AUC-ROC = 0.630, AUC-PR = 0.696) with a balanced operating point (sensitivity = 0.714, specificity = 0.435). This is appropriate when flagged compounds are ranked and forwarded for confirmatory testing rather than treated as definitive classifications.

An initial application of our approach to the compounds on the EDList has allowed us to identify a shortlist of compounds to evaluate in greater detail. Examples include clofentezine, 3-methylpyrazole, resorcinol, BHA, and thiabendazole, which combine the strongest MC-like scores with mechanisms plausibly detectable in a receptor-deficient line. Clofentezine, a known rodent hepatocarcinogen, received by far the highest score of the screen and is the strongest single candidate. 4-MBC also scored positive despite a predominantly estrogenic mechanism; since a receptor-mediated effect is not expected to be visible in U2OS, this may indicate a secondary, non-receptor activity and warrants cautious follow-up. In contrast, the three parabens all scored close to the decision boundary (propylparaben +0.009, methylparaben -0.002, butylparaben -0.008), indicating low-confidence classifications rather than a paraben class effect, as expected for compounds whose principal activity is receptor-mediated and therefore weakly resolved in U2OS. This process can be directly implemented by industries that need to anticipate chemical risks before conducting large-scale *in vivo* testing, particularly the food, pharmaceutical, and plastics industries, where it can help screen numerous compounds and focus experimental resources on those most at risk.

Some issues should be considered in a further study. The available data on cell morphology labeled as carcinogenic or non-carcinogenic in breast cancer remain limited in scope. In fact, there are 51 compounds remaining after excluding non-MC genotoxic compounds, which limits the reliability of covariance-dependent measures, such as Mahalanobis similarity. The Cell Painting profiles are also high-dimensional and noisy relative to this sample size: with 748 features for 51 compounds, similarity and distance calculations are prone to noise accumulation, which does not constitute an optimal framework for a small-sample dataset. This is why poor performance is observed in the full feature space, whereas dimension reduction (PCA) proved necessary to recover a usable signal. Furthermore, the profiles come from a single concentration (10 µM) and a single exposure duration (48 h), and a source-related batch effect persists despite Harmony correction (Korsunsky et al., 2019). Since the compounds were characterized and classified based on their ability to induce breast tumors in rodents, an additional *in vitro–in vivo* extrapolation gap separates the morphological score defined in our study from the actual risk to humans. The main concern, however, remains the cell line itself: the profiles were generated exclusively from U2OS osteosarcoma cells, which do not functionally express estrogen, progesterone, or androgen receptors. Consequently, when using JUMP-CP data, we cannot account for receptor-mediated endocrine mechanisms in the induced phenotypic response. It is therefore impossible to distinguish a “non-MC-like” score from a true absence of carcinogenic potential for compounds acting through this pathway, which constitutes a significant limitation, given that hormonal activity is a recognized pathway of breast carcinogenesis. A further consideration concerns the origin of the underlying data. The JUMP-CP profiles we reused were generated with U2OS cells cultured under standard conditions that, as is typical for this cell line, rely on fetal bovine serum (FBS), an animal-derived supplement obtained from fetal calf blood. Although this material was not used by us directly, since our study is entirely computational and reuses existing images, its presence in the source data means the resource is not fully animal-free. Future large-scale Cell Painting efforts intended to support NAM-based risk assessment would strengthen their 3R value by adopting serum-free or chemically defined media, or human platelet lysate, where compatible with assay performance.

This limitation related to the cell line clearly points to the most obvious perspective: replicating a similar analysis but using hormone-sensitive cell lines, such as MCF-7 or T47D, which would allow us to elucidate receptor-mediated endocrine mechanisms and interpret “non-MC-like” scores rather than dismissing them. Combining the U2OS cell line with a hormone-sensitive cell line in a single screening assay would allow us to address complementary mechanistic aspects. Another perspective would be to combine these phenotypic data with chemical structure data and/or transcriptomic data to gain a better understanding of the perturbations occurring within the cell.

## 5. Conclusion

Mammary carcinogenicity does not exhibit a single, specific phenotypic signature in U2OS cells. Instead, similar Cell Painting profiles cluster according to their mechanism of action. Genotoxic compounds produce strong, reproducible signals across different sources, whereas compounds with hormonal activity remain phenotypically silent, as the cell line used in JUMP-CP lacks functional steroid receptors. This heterogeneity limits overall supervised classification but aligns well with a Guilt-By-Association approach. In the best configuration identified of this approach which is based on an unsupervised PCA-reduced feature space, a cluster-centroid strategy with cosine similarity, we achieved an AUC-ROC of 0.630 and an AUC-PR of 0.696. Furthermore, the interpretable nature of our approach, which does not require training, enables broader applications to decipher certain underlying biological mechanisms. When applied prospectively to known and suspected endocrine disruptors, this framework identifies clofentezine, 3-methylpyrazole, resorcinol, BHA, and thiabendazole as candidates for confirmatory testing. These findings establish Cell Painting similarity as a usable, transparent triage tool for mammary carcinogenicity while making its boundaries explicit: a Non-MC-like score in an endocrine receptors-deficient cell line cannot rule out endocrine receptor-mediated risk. Validating flagged compounds in hormone-sensitive models and extending the screen to the broader range of endocrine-disruptor chemicals are the direct next steps toward a systematic, interpretable prioritization pipeline for environmental chemical risk assessment.

## Acknowledgements

We gratefully acknowledge the Fondation pour la Recherche Médicale (FRM) for its financial support of this research project [Grant ECO202306017371].

## Data availability

This study reused publicly available data. The mammary carcinogenicity annotations were obtained from the compendium of Kay and Rudel (2024). Cell Painting morphological profiles were obtained from the JUMP-CP dataset (cpg0016; https://github.com/jump-cellpainting/datasets) and processed with the jump-profiling-recipe workflow (https://github.com/broadinstitute/jump-profiling-recipe). Endocrine-disruptor candidate lists were retrieved from the EDLists database (https://edlists.org). All code used to compute morphological similarities, aggregate risk scores, and reproduce the analyses and figures in this study is available at https://github.com/rachebouche/mammary-carcinogenicity-cellpainting (archived at Zenodo, doi: 10.5281/zenodo.21512685, version [v1.0.0]). No new experimental data were generated in this study.

