## Supplementary figures and images for "A Cell Painting-Based Tool for the Risk Assessment of Mammary Carcinogens and Endocrine Disruptors"

### Supplementary Figure 1

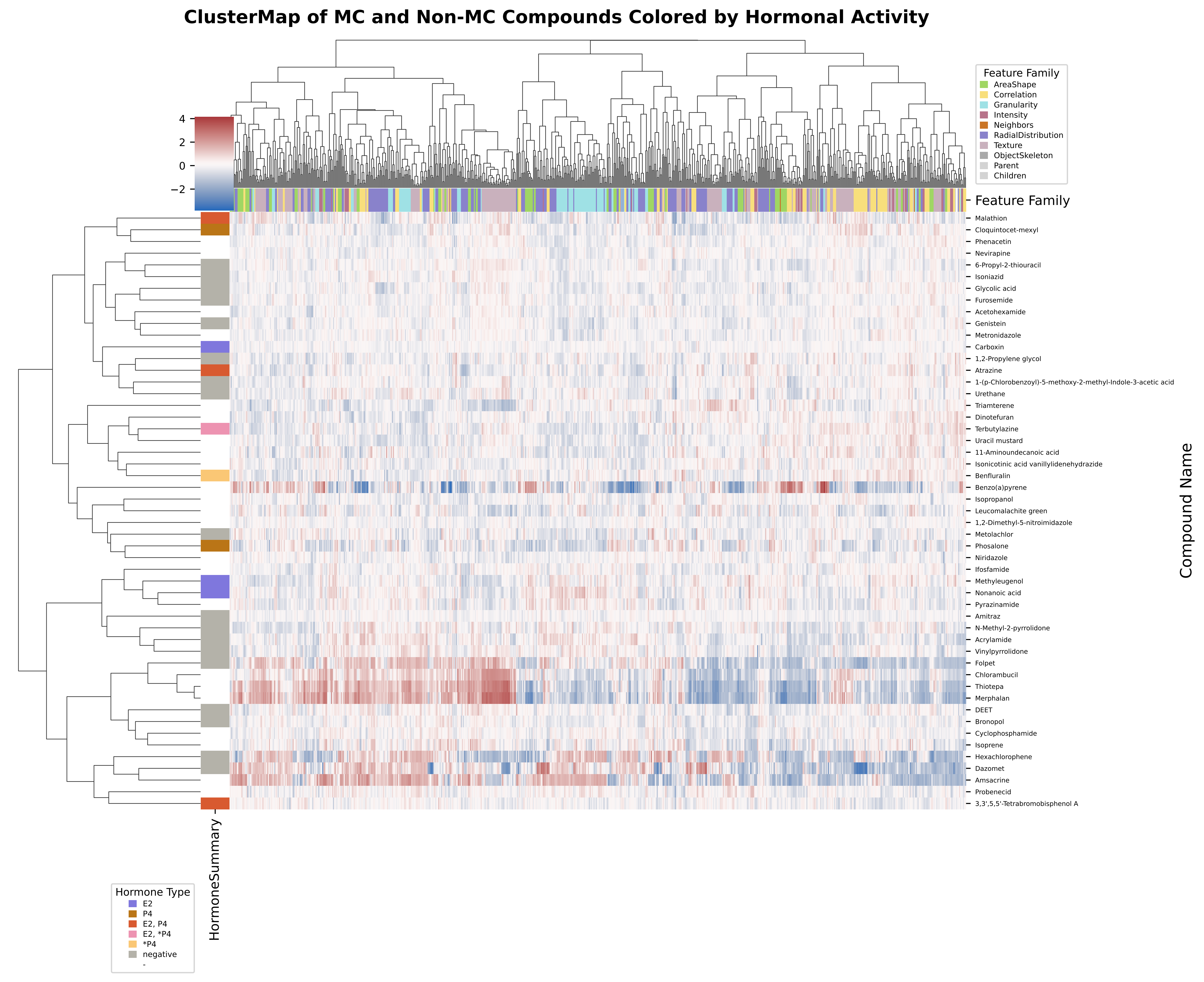
